# *Smad3* Regulates Smooth Muscle Cell Fate and Governs Adverse Remodeling and Calcification of Atherosclerotic Plaque

**DOI:** 10.1101/2020.09.15.299131

**Authors:** Paul Cheng, Robert C. Wirka, Juyong Brian Kim, Trieu Nguyen, Ramendra Kundu, Quanyi Zhao, Albert Pedroza, Manabu Nagao, Dharini Iyer, Michael P. Fischbein, Thomas Quertermous

## Abstract

Atherosclerotic plaques consist mostly of smooth muscle cells (SMC), and genes that influence SMC biology can modulate coronary artery disease (CAD) risk. Allelic variation at 15q22.33 has been identified by genome-wide association studies to modify the risk of CAD, and is associated with expression of *SMAD3* in SMC, but the mechanism by which this gene modifies CAD risk remains poorly understood. SMC-specific deletion of *Smad3* in a murine atherosclerosis model resulted in greater plaque burden, positive remodeling, and increased vascular calcification. Single-cell transcriptomic analyses revealed that loss of *Smad3* altered SMC progeny phenotype toward the previously described chondromyocyte fate, but importantly also promoted transition to a novel cell-state that governs remodeling and recruitment of inflammatory cells. This new remodeling population was marked by uniquely high *Mmp3* and *Cxcl12* expression, and its appearance correlated with higher-risk plaque features such as increased positive remodeling and macrophage content. Further, investigation of transcriptional mechanisms by which Smad3 alters SMC cell-fate revealed novel roles for Hox and Sox transcription factors whose direct interaction with Smad3 regulate an extensive transcriptional program balancing remodeling and vascular ECM with significant implications for human Mendelian aortic aneurysmal diseases. Together, these data suggest that *Smad3* expression in SMC inhibits the emergence of specific SMC phenotypic transition cells that mediate adverse plaque features, including positive remodeling, monocyte recruitment, and vascular calcification.

## Introduction

Decades of research and drug development have led to therapies and interventions that have significantly diminished morbidity and mortality from cardiovascular disease(*1, 2*). However, coronary artery disease (CAD) remains a leading cause of death in this country and worldwide with a sharp decline in the rate of improvement in mortality observed over the past decade(*3*). Recent large studies targeting well-characterized risk factors such as lipids (*4, 5*)and novel targets related to inflammation(*6, 7*) have had modest results(*8*) suggesting a continued need to identify new disease modifiers. Over the past decade, genome wide association studies (GWAS) have identified over 160 loci that contribute to CAD risk(*9, 10*). Causal variation identified in these loci point primarily to genes and pathways predicted to function in the blood vessel wall to regulate disease risk (*11-13*).

Specific features of atherosclerotic plaques have been increasingly recognized to offer significant prognostic value. For instance, it has been noted that cellular composition, such as SMC contribution to the fibrous cap, influences risk of plaque rupture(*14-16*). With advances in diagnostic imaging, novel features such as positive remodeling and microcalcification have been found to be highly predictive for myocardial infarctions(*17, 18*). Positive remodeling, defined as expansion of the vascular wall beyond its original confines without narrowing the vascular lumen, is thought to represent sites at risk for impending plaque rupture(*17-20*). However, little is known regarding the cellular and molecular mechanisms by which these high-risk features are controlled.

A number of post-genomic GWAS follow-up studies in this and other laboratories investigating the genetic disease-related mechanisms of CAD have focused on SMC, and the relationship of SMC cell state changes to disease risk(*21-25*). These studies have indicated that a significant portion of the CAD attributable risk is determined by this cell type(*13*). Consistent with this notion, recent lineage tracing studies have demonstrated that the majority of cells inside atherosclerotic plaque, including those expressing some inflammatory markers, are oligo-clonal de-differentiated smooth muscle derivatives(*26-29*). Genes that alter SMC behavior are known to influence the composition of an atherosclerotic plaque(*21, 24, 25, 28, 30*). CAD-associated genes *TCF21* and *AHR* have been linked to these processes, and various genomic data suggests that additional CAD genes might also regulate SMC phenotype(*23, 30-34*). While these SMC progenies have been previously lumped together as phenotypically modulated SMC, advances in single cell RNA profiling has demonstrated the presence of subsets of these cells with distinct transcriptomes and cell fates. For example, medial SMC have been shown to give rise to fibroblast-like cells termed fibromyocytes(*25*), as well as cells similar to endochondral bone forming cells, chondromyocytes(*21, 32*), among other bioinformatically defined populations(*21, 35*). However, the functional significance and relative location of these transcriptionally distinct populations remain to be elucidated.

The TGFβ signaling pathway is central to smooth muscle biology during development and disease, and an important modifier of atherosclerosis(*36, 37*). Canonical TGFβ signaling is thought to be mediated through Smad family proteins, particularly nuclear signaling factors Smad2 and Smad3. While themselves poor binders to DNA(*38*), through their interaction with other transcription factors, the Smad factors are central to key transcriptional programs that regulate cell fate in development and disease(*39, 40*). Multiple GWAS have identified rs17293632(*9*), a single nucleotide variant that lies within a functional smooth muscle enhancer that regulates SMAD3 expression(*22*), as an important modifier of risk for myocardial infarction. However, how smooth muscle *SMAD3* expression influences risk of myocardial infarction is unclear. *In vitro*, SMAD3 appears to modify smooth muscle cell differentiation and proliferation through its interaction with other transcription factors critical to SMC biology and risk of CAD(*31*). However, the exact effects of *Smad3* expression level on SMC plaque biology remain unknown.

Here, we demonstrate that SMC-specific deletion of *Smad3* influences the fate of de-differentiated SMC in atherosclerotic plaques *in vivo*, promoting both a new pro-remodeling SMC transition phenotype that expresses remodeling genes such as *Mmp3* and inflammatory chemokines such as *Cxcl12, as well as* an expansion of SMC-derived chondromyocyte (CMC) population. These cellular changes are associated with increased positive remodeling and plaque calcification that appear to be directed by Smad3 in conjunction with transcriptional effects of Hox and Sox factors.

## Results

### SMC-specific deletion of Smad3 is associated with increased lesion burden, outward remodeling, and increased numbers of both SMC progeny and monocyte/macrophage lineage cells

To understand how smooth muscle expression of *Smad3* influences atherosclerotic lesions *in vivo*, we generated a murine model of atherosclerosis with established smooth-muscle specific Cre (*Myh11-Cre*) crossed with a conditional knockout allele of *Smad3*(*41-43*) with concurrent lineage tracing (*ROSA*^*Tdt*^) on the *ApoE* null background (*Smad3*^*ΔSMC*^) (Fig. 1A). To limit confounding created by the critical role of Smad3 during development, *Smad3* was deleted via a tamoxifen inducible Cre only after mice had reached maturity (8-weeks-old), immediately prior to initiation of a Western high-fat diet (HFD, Fig. 1A). The *Smad3* conditional knockout mice grew to maturity with no significant change in weight or mortality compared to control (Suppl. Fig. 1A, B), with highly efficient Cre-mediated deletion of the floxed region of *Smad3*, i.e., the DNA binding domain) (Suppl. Fig 1C).

**Figure 1:**
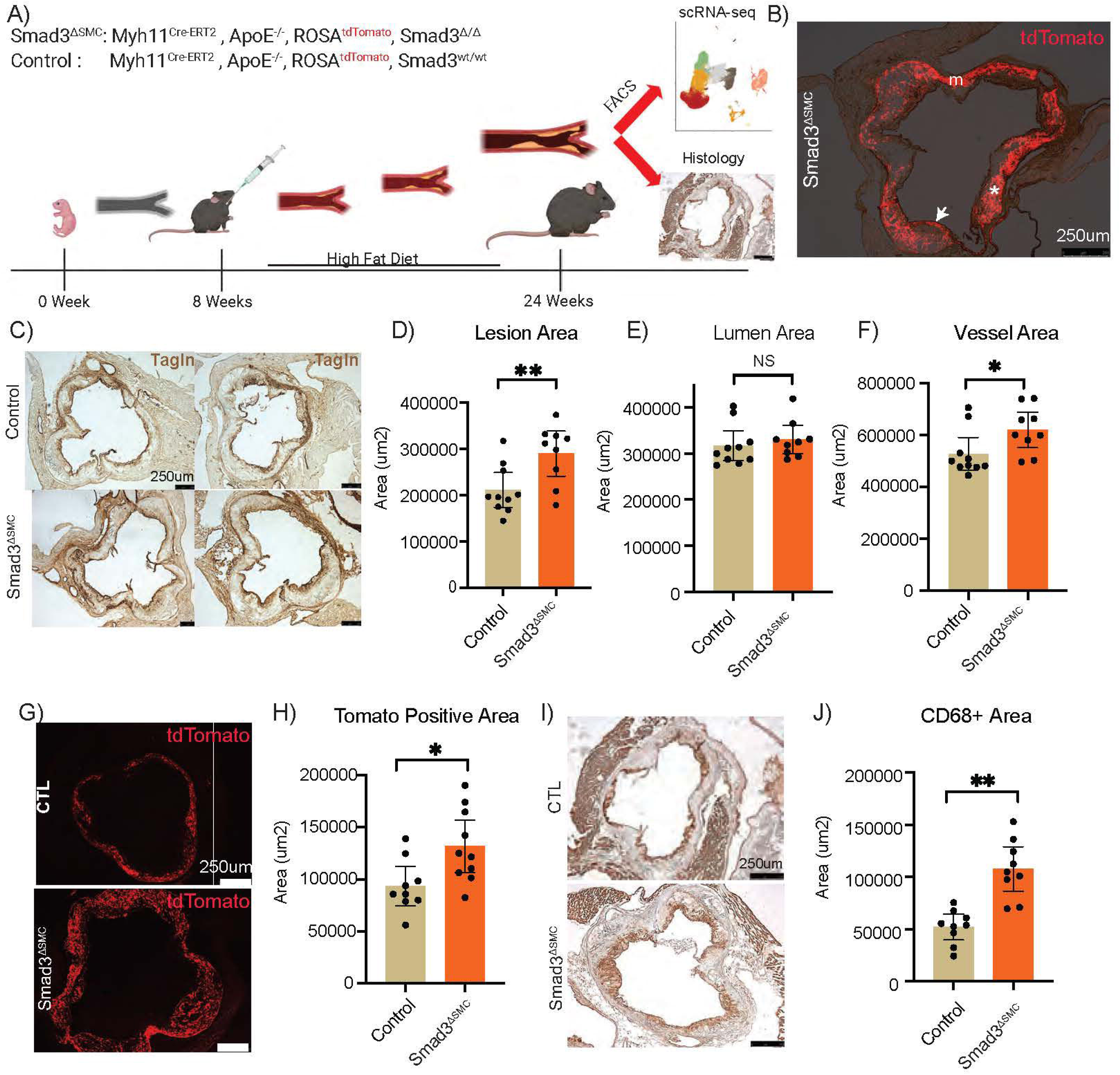
*Smad3*^*ΔSMC*^ mice have increased lesion burden in the *ApoE* null model. (A) Representative figure of mouse protocol showing SMC-specific lineage tracing and Smad3 conditional knockout (*Smad3*^*ΔSMC*^). (B) *Smad3*^*ΔSMC*^ SMC transition cells can contribute to all regions of the lesion, including the lesion cap (arrow), modulated SMC (*), and tunica media (m). (C) Representative sections from control and *Smad3*^*ΔSMC*^ mice stained for Tagln to highlight cap and tunica media for lesion quantification. (D) Lesion area between the lumen and internal elastic lamina, (E) lumen area, (F) vessel area quantified as area between lumen and external elastic lamina, across experimental animals. (G) Representative sections from control and *Smad3*^*ΔSMC*^ mice revealing tdT fluorescence to show lineage traced cells in the media and plaque. (H) Quantification of tdT positive area in control and *Smad3*^*ΔSMC*^ mice. (I) Representative sections from control and *Smad3*^*ΔSMC*^ mice stained with CD68 antibody. (J) Quantification of CD68 positive area,* p<0.05. Each dot from each bar graph represents data measured from a single matched section from each individual animal. Error bars represent 95% CI of mean.

Examination of atherosclerotic lesions in the aortic root after 16 weeks of HFD demonstrated that SMCs from *Smad3*^*ΔSMC*^ mice were able to migrate into the lesion, expand, and contribute to the formation of atherosclerotic plaque and the fibrous cap (Figs. 1B, 1C). Quantification of atherosclerotic lesions revealed a significant increase in plaque volume in *Smad3*^*ΔSMC*^ compared to control animals (Fig. 1D). To further characterize the anatomy of diseased vessels, specifically regarding outward remodeling vs luminal narrowing, we evaluated the area encapsulated by the diseased vessel as well as lumen area. The lumen area in these sections showed no significant change (Fig. 1E), but the area circumscribed by the external elastic lamina was significantly increased (Fig. 1F). These findings are consistent with expansion of atherosclerotic plaque volume in conjunction with outward “positive” remodeling. To determine the cellular anatomy associated with the increased plaque size, we quantified the area occupied by fluorescent tdTomato SMC lineage traced cells (Fig. 1G, 1H) as well as CD68 stained monocytes and macrophages in the lesions (Fig. 1I, 1J). This analysis revealed a statistically significant increase in area for both SMC-derived cells as well as cells of the monocyte-macrophage lineage. Given that the lineage tracing *Myh11-Cre* transgene is active only in cells emanating from mature SMC, these findings suggest that the increased lesion growth has both a cell autonomous as well as a non-autonomous cellular component, with the latter reflecting an SMC mediated effect on monocyte-macrophage lesion recruitment.

### Single cell transcriptomic profiling identifies an expansion of SMC-derived chondromyocyte phenotype cells, and a novel SMC lineage cell transition phenotype, with Smad3 deletion

Given the critical role that *Smad3* plays in cell-fate decisions during development, we hypothesized that alteration in disease-associated-SMC phenotype transitions might account for the observed cellular lesion characteristics as well as recruitment of CD68+ cells and positive remodeling. Thus, to better understand how loss of *Smad3* expression produced phenotypic changes in lesion SMC derived cells, and their interactions with other lesion cell types, we performed single cell RNA expression profiling (scRNAseq) of atherosclerotic lesions from *Smad3* knockout and control animals. The atherosclerotic tissue was harvested, processed for single cell encapsulation, RNA capture, reverse transcription and amplification with the 10X Genomics Chromium V3 platform, and cDNA libraries were sequenced as previously described(*25, 30*). Subsequently, the single cell expression data were visualized utilizing Uniform Manifold Approximation and Projection (UMAP) to create a 2-D projection representing the organization of individual cells to each other in clusters, and of the relationship of the organized clusters to each other. After filtering and normalization, a total of 26,219 cells from control and 35,518 cells from *Smad3*^*ΔSMC*^ mice were included in the analysis obtained from 3 and 4 independent captures of pooled tissue from 2 mice in each individual capture.

We first investigated clustering of the scRNAseq data using standard settings for the principal component analysis and number of clusters in UMAP space. Feature plots were employed to visualize the SMC lineage traced cells (Fig. 2A). SMC traced cells were identified by expression of the *tdTomato* gene, and the component of this cluster representing mature medial SMC was identified by expression of SMC lineage markers *Myh11* and *Cnn1*. As we have shown previously, a significant portion of the SMC-lineage traced cells during disease did not express these mature SMC markers and thus represented SMC that had undergone phenotypic transition(*24, 25, 30*). We were thus able to determine whether there were alterations in the number of mature differentiated versus transition SMC in *Smad3*^*ΔSMC*^ mice. We investigated whether there was an alteration in the number of differentiated SMC among the lineage traced (tdTomato+) cells, defining “differentiated” cells using classical Myh11 or Cnn1 expression, along with unbiased clustering (SMC vs transition SMC) (Fig. 2A, 2B). Cells were clustered based on the scRNAseq data using the Lovain algorithm to a resolution where the SMC derived cells are split into two distinct clusters, as we and other have characterized(*25*).

**Figure 2:**
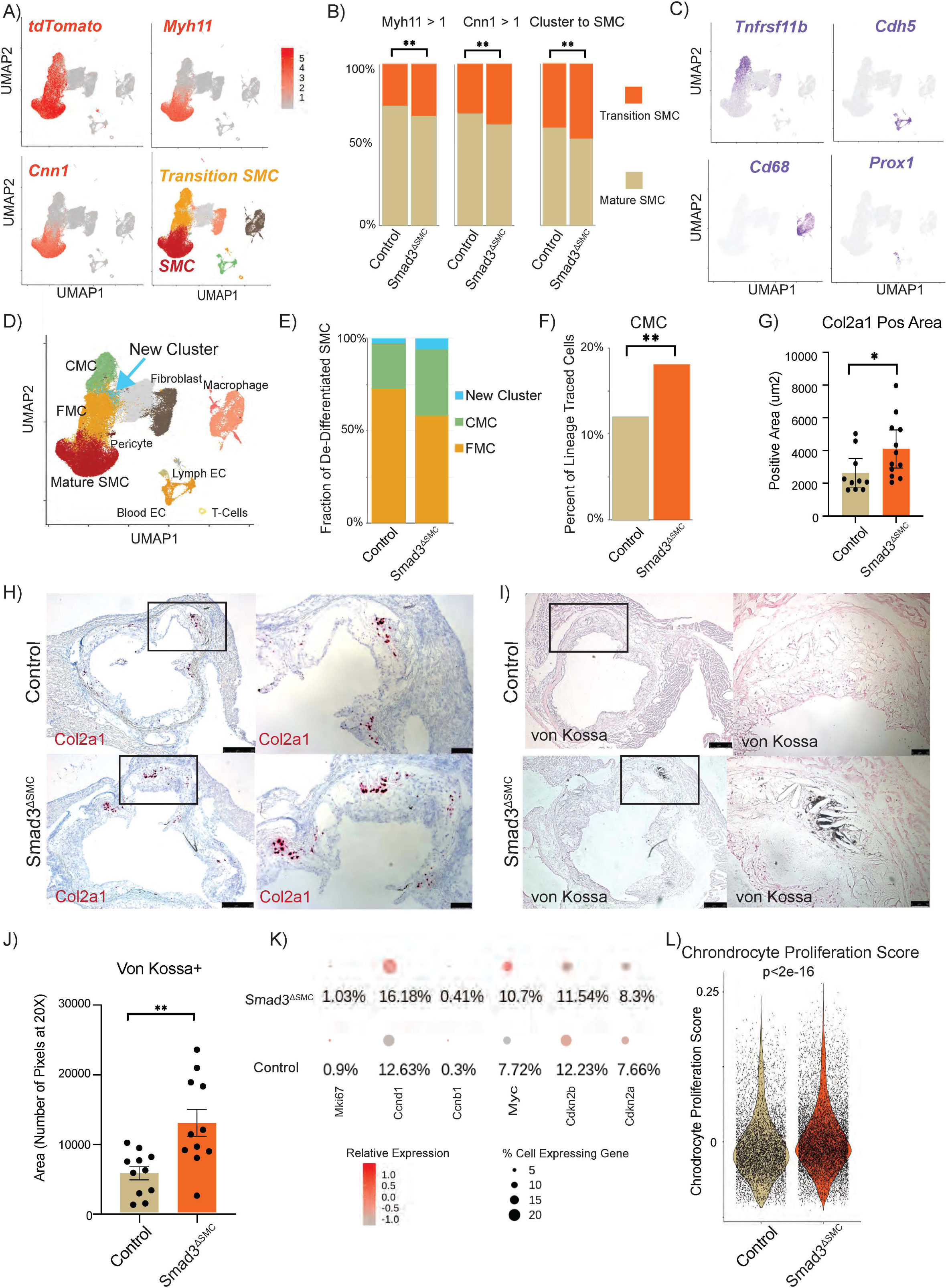
Loss of *Smad3* alters cell-fate decisions of transition SMC in atherosclerotic lesions. (A) Expression of lineage tracing marker tdTomato, mature SMC markers *Myh11, Cnn1*, and unbiased clustering identified transition SMC in UMAP depiction of scRNAseq data. (B) Fraction of lineage traced cells that were mature SMC as defined by *Myh11, Cnn1* expression or unbiased clustering. (C) Expression of key lineage markers of other disease relevant cells captured by our scRNA sequencing, including fibromyocytes (*Tnfrsf11b*), endothelial cells (*Cdh5*), macrophages (*CD68*), and lymphatic endothelial cells (*Prox1*). (D) Unbiased clustering of all captured cells and respective clustering upon applying the Louvain algorithm, represented in UMAP space, with their respective biological identities as defined by marker genes. (E) Fraction of de-differentiated lineage-traced cells that contribute to fibromyocytes (FMC), chondromyocytes (CMC), and the newly identified cell cluster in control and *Smad3*^*ΔSMC*^ mice. (F) Fraction of lineage traced cells that contribute to CMC in control and *Smad3*^*ΔSMC*^ mice. (G) CMC marker *Col2a1* expression by RNAscope, positive area per section in control and *Smad3*^*ΔSMC*^ mice. (H) Representative images of colorimetric *Col2a1* RNAscope quantification with control and *Smad3*^*ΔSMC*^ mice. (I) Von Kossa staining of sections from control and *Smad3*^*ΔSMC*^ mice, with (J) quantification as area of staining. (K) Dot plot representation of expression of cell-cycle markers in transitional SMC as represented by relative expression (color) and fraction of total positive cells (size). (L) Chondrocyte proliferation score of control and *Smad3*^*ΔSMC*^ transitional SMC. * p<0.05, ** p<0.01.

Regardless of the definition of “mature SMC”, there was a significant decrease in the proportion of mature SMC in the *Smad3*^*ΔSMC*^ compared to control mice (Fig. 2A,B).

To discern the optimum biologically relevant resolution of clustering and avoid over-fitting with the Lovain algorithm, the clustering resolution was empirically determined as the minimal settings that allowed for separation of known biologically distinct populations of endothelial cells (EC) (Figs. 2C, D), i.e., blood vessel EC (VE-cadherin expressing, Prox1 negative) vs lymphatic EC (Prox1 expressing) endothelial cells(*44, 45*). The identities of the cellular subpopulations that were created with this approach were then determined based on specifically expressed genes in each cellular cluster with the monocyte-macrophage lineage identified with CD68 expression, and the previously described SMC transition fibromyocytes by the marker *Tnfrsf11b* (Fig. 2C). The validity of this clustering approach was further confirmed via visualization of cell population canonical “markers” for all clusters in the UMAP space. Previous work in SMC lineage tracing in atherosclerosis has identified three distinct clusters of SMC derived cells, including mature SMC, fibromyocytes (FMC), and pro-calcific chondromyocytes (CMC)(*21, 30, 35, 46*). Similar subsets of lineage traced SMC were identified in these data (Fig. 2D), and in addition two distinct new clusters of Myh11-Cre lineage labeled populations were also identified. One cluster was composed of pericytes, and an additional small population of novel disease-associated SMC-derived cells clustered by itself as a unique transcriptomic phenotype.

To determine if loss of *Smad3* expression alters the cell-fate transitions of disease-associated SMC lineage cells, we ascertained the fraction of de-differentiated cells that contribute to clusters for each of the two known and the novel transition phenotypes. *Smad3*^*ΔSMC*^ transition SMC showed an increase in proportion of CMC along with the newly defined population at the expense of the more differentiated FMC (Fig. 2E, F). The increase in fraction of CMC among disease-associated *Smad3*^*ΔSMC*^ cells corresponded with a significant increase in lesion area expressing chondromyocyte marker *Col2a1* (Fig. 2G, H). Functionally, this increase in *Col2a1* expressing cells correlated with an increase in lesion calcified area as assayed by von Kossa staining of atherosclerotic lesions (Fig. 2I, J). These cells appeared to result from an increase in proliferation of de-differentiated SMC, as evidenced by a higher level and increased proportion of cells expressing proliferation markers such as *Mki67, Ccnd1, Ccnb1, Myc, and Cdkn2a/b* (Fig. 2K). To further assess this possibility, we generated a “chondrocyte proliferation score” that represents a scaled-average expression of all genes associated with GO category “promote chondrocyte proliferation” with each individual cells’ transcriptome(*47*). Consistent with higher expression of specific cell-cycle regulators, *Smad3*^*ΔSMC*^ transition SMC had a significantly higher chondrocyte proliferation score than control cells (Fig. 2L). Applying the analysis utilizing a “mesenchymal proliferation score” based on GO categories of “promote mesenchymal proliferation” produced equally significant results (Sup. Fig. 2), suggesting that the increase in number of tdTomato positive cells in the lesions likely reflects proliferation of SMC derivative cells. The increase in cell number did not result from alterations in apoptosis, since there was no difference in TUNEL staining of control vs *Smad3*^*ΔSMC*^ sections (Sup. Fig, 3).

### A novel SMC transition cell phenotype is characterized by expression of Mmp3 and leukocyte recruiting factors

In addition to increased CMC, there was also an increase in the proportion of SMC-derivatives that contributed to the newly identified SMC transition cell cluster (Figs. 2E, 3A), that we will refer to as remodeling-SMC (R-SMC) due to their high expression of genes involved in extracellular matrix remodeling. The vast majority of cells within this cluster were lineage traced at a proportion similar to that seen with CMC and FMC (Fig. 3B) confirming they were SMC derived. In vascular lesions, they constituted 3% of all lineage traced cells and 6% of SMC transition cells in *Smad3*^*ΔSMC*^. While their gene expression pattern is most similar to that identified in CMC as shown by their juxtaposition in UMAP space and by hierarchical clustering, they were easily distinguished from CMC at the transcriptional level (Fig. 3D). To investigate the ontogenic relationship of this group of cells to SMC, FMC, and CMC, we ported the scRNAseq data to Slingshot(*48*), a lineage inference tool designed to map trajectories involving multiple branching lineages. Applying this algorithm suggested that SMC give rise to FMC which in turn serve as a source for both CMC and R-SMC (Fig. 3C).

**Figure 3:**
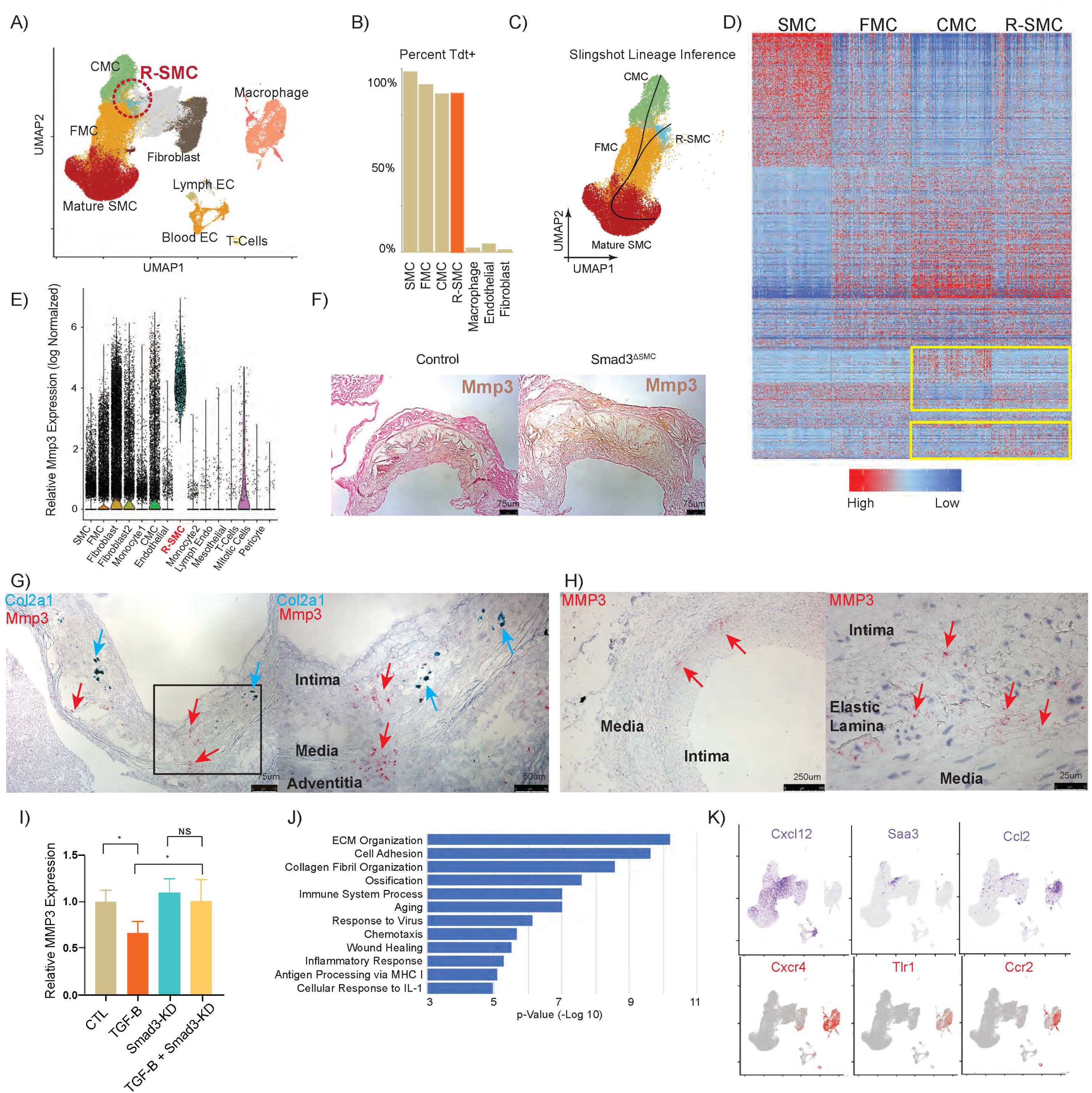
The novel cluster of *Mmp3*-expressing R-SMC transition cells promote remodeling and inflammation. A) UMAP representation and location of the novel cluster of cells with high *Mmp3* expression. (B) Percent of tdTomato positive cells representing different cellular clusters. (C) Pseudotemporal alignment of cells with Slingshot lineage inference (solid black line) identified a likely relationship for SMC, FMC, CMC, and R-SMC. (D) Heatmap of differentially regulated genes among distinct lineage-traced SMC in the lesion. Differentially expressed genes between CMC and R-SMC are indicated by yellow boxes. (E) Normalized *Mmp3* expression of all the cells in the lesion. (F) Immunohistochemistry of Mmp3 in control and *Smad3*^*ΔSMC*^ mice. (G) RNAscope of *Mmp3* (red) and *Col2a1* (blue) expressing cells in atherosclerotic lesion showing distinct localization and clustering. (H) RNAscope for *MMP3* expression in human coronary artery speciments. (I) Relative *MMP3* expression in HCASMC in the presence and absence of TGFβ and SMAD3. (J) Biological processes enriched in genes that are preferentially expressed by R-SMC compared to other transitional SMC. (K) Expression of chemoattractants (Cxcl12, Saa3, Ccl2) that are specific to R-SMC with their respective receptors (Cxcr4, Tlr1, Ccr2) showing expression restricted to the monocyte cluster, * p<0.05. Each dot from each bar graph represents a single qPCR measurement from each biological replicate. Error bars represent SEM.

Interestingly, the R-SMC cell population was marked by a particularly high expression of matrix metalloproteinase-3 (*Mmp3*) (Figs. 3E, 3F), an enzyme required for outward remodeling of atherosclerotic vessels(*49*). Given our observation of increased outward remodeling identified in *Smad3*^*ΔSMC*^ mice, this finding was further investigated. Single-cell transcriptomic data suggested a higher number of *Mmp3*-expressing cells and an overall higher level of *Mmp3* expression in *Smad3*^*ΔSMC*^ than control cells (Sup. Fig. 4A, 4B, 4E). *In situ* hybridization with RNAscope showed *Mmp3*-expressing R-SMCs to be a distinct population from CMC, which were marked by *Col2a1* expression (Fig. 3G), indicating that these R-SMC transition cells are juxtaposed but not overlapping in the lesion plaque and further supporting their distinct phenotype. Consistent with the established role for MMP3 in positive remodeling, its expression was most prominent at the base of the atherosclerotic lesion in cells juxtaposed to the elastic lamina. Strikingly, these MMP3 cells were consistently associated with areas of disrupted elastic lamina, consistent with their invasion through this structure (Fig. 3F, 3G, Sup Fig 4D).

**Figure 4:**
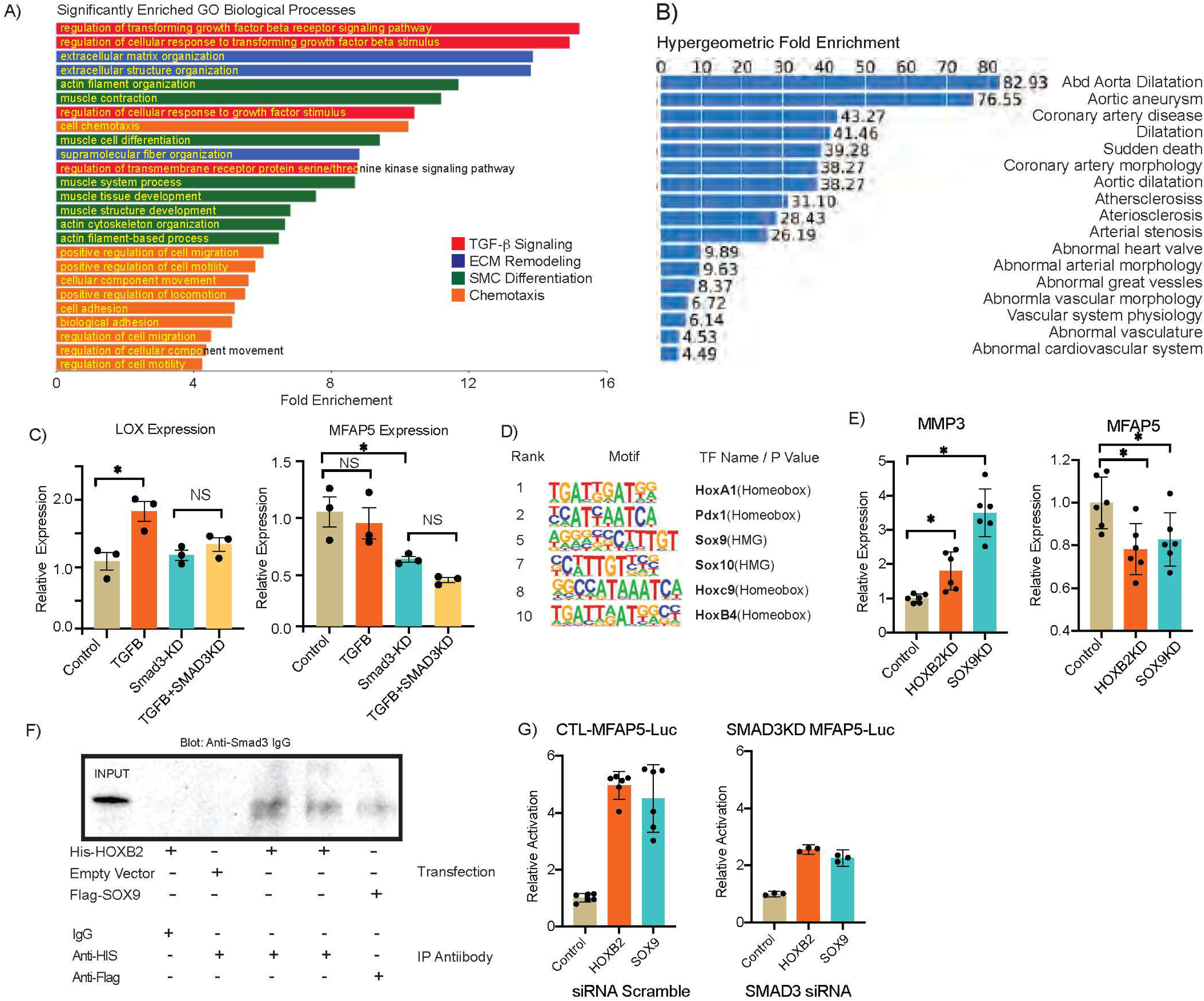
Smad3 regulates a transcriptional program associated with vascular outward expansion in conjunction with Sox9 and HoxB2. (A) Significantly enriched GO biological processes represented by Smad3 differentially regulated genes. (B) Enrichment of human disease terms associated with differentially regulated genes. (C) Relative expression of *LOX* and *MFAP5* in HCASMC in the presence and absence of TGFβ, in cells transfected with siRNA against *SMAD3* (SMAD3KD) or non-targeting control. (D) HOMER analysis results for motifs enriched in promoter regions of genes differentially regulated in *Smad3*^*ΔSMC*^ mice. (E) Expression of *MMP3* and *MFAP5* in control and *SOX9* or *HOXB2* knockdown HCASMC. (F) Co-immunoprecipitation of His-tagged HOXB2 (lane 4/5) or Flag-tagged SOX9 (lane 6) with endogenous SMAD3 compared to IgG or control vector (lanes 2/3). (G) Relative luciferase activity of MFAP5-Luc in HEK-293 cells transfected with HOXB2 and SOX9 expression vectors in the absence (left) or presence (right) of SMAD3 siRNA (SMAD3KD) or control siRNA (control), *p<0.05. Each dot from each bar graph represents a single normalized luciferase measurement from each biological replicate. Error bars represent SEM.

Importantly, the same rare population of MMP3 expressing cells were also found in human coronary arteries, where they were also associated with regions of disrupted elastic lamina (Fig 3H). There were more cells expressing Mmp3 in *Smad3*^*ΔSMC*^ compared to controls as per the scRNAseq data and more prominent staining in *Smad3*^*ΔSMC*^ vs control (Fig. 3F, Sup Fig 4A). *Mmp3* expression was altered only in the SMC derived cells, and not significantly different in the non-SMC derived population, suggesting that altered Mmp3 levels by Smad3 happens in a cell-autonomous manner in SMC lineage transition cells (Sup Fig 4a). By performing TGF-β stimulation of human coronary artery smooth muscle cells, we were able to show that MMP3 is suppressed by TGF-β (Fig. 3I). SMAD3 knockdown did not significantly increase the expression of MMP3 at baseline, but did negate the TGF-β dependent suppression of expression, suggesting SMAD3 is required for the TGF-β dependent regulation of MMP3 in atherosclerotic lesions.

To better understand the vascular function of this cell population, we performed pathway analyses of all significantly differentially expressed genes in these cells as compared to other de-differentiated SMC phenotypes (Sup Table 1). PANTHER analysis revealed the top biological processes enriched in this list of genes are related to remodeling of ECM followed by regulation of chemotaxis and inflammation (Fig. 3J). Analysis of genes contributing to the enrichment of these pathways reveal that this population of cells express higher levels of multiple chemoattractants whose receptors are primarily restricted to macrophages (Fig. 3K). For example, this population is the main SMC source of Cxcl12 in the lesion, with the Cxcr4 receptor for this ligand being expressed exclusively by lesion macrophages. In addition, R-SMC differentially expressed additional cytokine genes, including *Saa3*, and *Ccl2*, which all have well-established roles in monocyte recruitment, with their respective receptor genes *Tlr1* and *Ccr2* expressed mostly by inflammatory cells in the lesion (Fig. 3K). Taken together, these findings suggest that beyond its role in positive remodeling, this novel population of cells may orchestrate monocyte recruitment and underlie the increase in CD68 positive cells in lesions.

### Smad3 cooperates with other transcription factors to regulate SMC genes that are involved in human vascular remodeling and disease

Given that *Smad3* is likely affecting all SMC lineage phenotypes beyond just the CMC and R-SMC populations described above, we characterized the alteration in gene expression of all de-differentiated SMC derived cells in the atherosclerotic lesions. To rigorously identify differentially regulated genes, we employed a Wilcoxon rank sum-based analysis comparing *Smad3*^*ΔSMC*^ versus control mouse data for FMC, CMC and R-SMC cellular clusters and 83 genes were identified (Supplmentary table 2). Pathway analysis performed using DAVID demonstrated significant enrichment in TGFβ signaling, which is consistent with Smad3’s central role in the TGFβ pathway (Fig. 4A). In addition, biological processes broadly related to ECM remodeling, SMC differentiation, and chemotaxis were also heavily enriched. Utilizing GREAT(*50*), we compared the list of 83 identified *Smad3*^*ΔSMC*^ marker genes to the top 1000 genes expressed by de-differentiated SMC in diseased murine vessels. The analysis revealed enrichment for genes linked to human atherosclerosis and arterial dilatation (Fig. 4B). In addition to *Mmp3*, and consistent with the observed phenotype of increased outward remodeling in *Smad3*^*ΔSMC*^ mice, *Lox* and *Mfap5* were also among the top significantly down-regulated genes in de-differentiated *Smad3*^*ΔSMC*^ SMC (Sup table 2). Pathogenic loss of function in both of these genes has been linked to human vascular syndromes including aortic aneurysms(*51-53*). To investigate whether these genes are directly regulated by TGFβ, we stimulated human coronary artery smooth muscle cells (HCASMC) with TGFβ in the presence and absence of *SMAD3*. A subset of genes, such as *LOX*, appear to be directly regulated by TGFβ in HCASMC, and this effect was abrogated by *SMAD3* silencing (Fig. 4C), similar to the regulation of *MMP3*. However, a subset of genes, such as *MFAP5*, did not appear to be TGFβ responsive, but were sensitive to loss of *SMAD3* (Fig. 4C). These findings suggest involvement of additional co-regulatory factors.

To better understand the context in which Smad3 regulates this complex transcriptional program and identify possible co-regulatory factors, we analyzed the 5’ regulatory elements of the 83 Smad3 differentially regulated genes to look for enriched transcription factor motifs. Motif analysis conducted with HOMER revealed enrichment for Hox/homeobox motifs as well as Sox9/10 related motifs (Fig. 4D). *Hox* and *Sox* genes were not down-regulated in the *Smad3*^*ΔSMC*^ SMC (Sup. Fig. 5), suggesting that these factors are likely directly interacting with Smad3 to regulate a joint transcriptional program. Consistent with this hypothesis, Sox9 is known to be a critical transcription factor regulating calcification, and has been shown in chondrosarcoma cells to selectively interact with Smad3, but not Smad2, to modulate an endochondral ossification program(*54*). Knocking down *SOX9* and *HOXB2*, the most highly expressed *HOX* gene in human coronary SMC, recapitulated the changes in expression of key vascular remodeling genes *MMP3* and *MFAP5* observed *in vivo* in mouse (Fig. 4E). Thus, we hypothesize that HOX factors, whose motifs are also enriched, can directly interact with SMAD3. To test this hypothesis, we performed nuclear Co-IP experiments to test their interaction. In cells over-expressing His-HOXB2 or Flag-SOX9, SMAD3 protein was co-immunoprecipitated with anti-His or Flag antibody respectively (Fig 4F, Sup Fig 6), suggesting HOX family proteins such as HOXB2 are physically bound to SMAD3 in the nucleus, as is SOX9. To test the functional significance and epistatic relationship of these findings, we cloned an evolutionarily conserved regulatory region near the 5’ end of the human MFAP5 gene, which contains conserved putative HOX, SOX, and SMAD binding elements, into a luciferase vector and tested its ability to respond to SOX9 and HOXB2 binding. SOX9 and HOXB2 efficiently activated this enhancer element but not the control luciferase vector, further establishing a role in this transcriptional program (Fig 4G, Sup Fig 5B). Knocking down SMAD3 diminished the SOX9 and HOXB2-dependent activation of the luciferase construct, suggesting interaction with SMAD3 is required for regulation of MFAP5 (Fig 4G).

**Figure 5:**
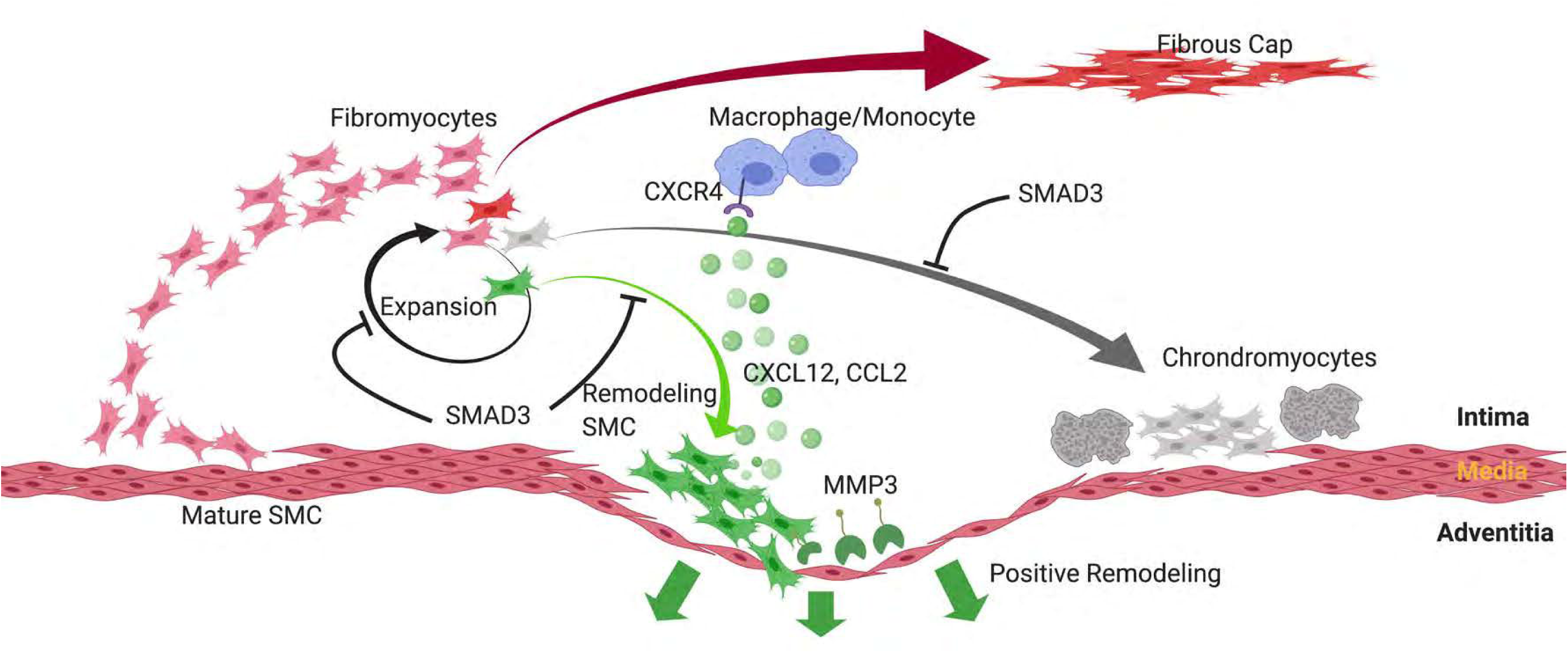
Smad3 regulates smooth muscle cell-fate and governs adverse remodeling and calcification of atherosclerotic plaque.

## Discussion

In these experiments we have studied *Smad3*^*ΔSMC*^ mice with scRNAseq, cellular anatomy, and lesion *in situ* studies. Highly correlated results from these different approaches provide a consistent picture of the role of this CAD associated transcription factor in vascular disease processes. Applying scRNAseq at unprecedented depth and cell number combined with lineage tracing allowed us to observe changes in the number of critical SMC transition cells and transcriptomic changes that were previously unobtainable. These data identified cell state changes related to loss of *Smad3* expression, such as increased SMC transition to the CMC phenotype, a finding substantiated with increased *Col2a1* plaque cell expression and vascular calcification. Furthermore, the loss of *Smad3* promoted SMC transition to a unique SMC phenotype cell, R-SMC, which mediates an apparent role in modulating multiple plaque characteristics. Increased expression of cellular proliferation genes by scRNAseq was consistent with the increased number of SMC lineage traced cells in the plaque (Fig. 5), echoing previous *in vitro* findings in HCASMC(*31*). Finally, leveraging rich transcriptomic data from a large number of cells, we identified transcriptional interaction of SMAD3 with HOX and SOX factors in the regulation of genes expressed in de-differentiated SMC transition phenotypes.

The high resolution scRNAseq has data allowed us to describe a small but distinct subset of SMC transition cells that expresses both pro-remodeling enzymes as well as pro-recruitment chemokines. Mmp3, a metalloprotease required for positive remodeling, is expressed most prominently in the R-SMC that are located in the basal plaque and appear to migrate into the media towards the adventitia, potentially related to the previously described SMC lineage traced population in the adventitia(*55*). Strikingly, in human coronary arteries, this MMP3+ pro-remodeling population resides in the same regions of the artery with observed disruption of the nearby elastic lamina, consistent with a likely role of promoting positive remodeling in human coronary disease. These findings suggest that expansion of this population may be a significant contributing factor to positive remodeling and other adverse features. The importance of this MMP3-expressing SMC transition population in modifying plaque rupture risk may explain the seemingly paradoxical observation that a SNP associated with lower-expression of MMP3 is associated with greater luminal coronary artery stenosis on cardiac catheterization, but the higher-expressing variant is associated with more myocardial infarctions(*56*). These genetic observations provide further evidence that modulation of R-SMC alters plaque features and CAD-risk in human. While *Mmp3* was previously thought to be expressed by macrophages in the lesions(*56*), we found no evidence that *Mmp3* was expressed by cells in the macrophage cluster. This was likely due to previous erroneous identification of SMC transition cells expressing inflammatory markers as macrophages(*25, 57*). Previous work has shown that IL1 drives positive remodeling of plaques and this function was completely reversed by deletion of *Mmp3*(*49*). This suggests that SMC are the primary mechanism for the high-risk plaque feature of positive remodeling, as seen in the *Smad3*^*ΔSMC*^ mice, and may account in part for the beneficial effect of IL1 blockade on the risk of plaque rupture in humans(*6*). *Mmp3* also appears differentially regulated among different population of SMC progenies in the carotid artery plaque(*21*), suggesting a similar R-SMC population may exist in other atherosclerotic beds. Interestingly, this same population of cells express a number of chemokines including *Cxcl12*, whose main receptor *Cxcr4* is expressed solely on the monocyte/macrophage lineage. This suggests that the R-SMC population likely plays an important role in regulating the inflammatory response to the lesion, contributing to the increase in observed monocyte/macrophage population detected in the lesions. The combination of remodeling and inflammatory cell recruitment, both critical factors of plaque stability, highlights the critical role that this specific sup-population of cells may play in modulating human disease risk. This contributes to existing literature(*58*) suggesting that SMC play a central role in regulating inflammatory cell recruitment and retention in atherosclerotic plaque and identifies a specific sub-population of transition SMC critical for high-risk plaque features.

There was also an increase in CMC in *Smad3*^*ΔSMC*^ mice suggesting Smad3 actively inhibits differentiation to this phenotype or inhibits their proliferation. The transcriptomic, topological, and lineage inference data presented here suggest they are a distinct population from *Mmp3*-expressing R-SMC. The CMC exhibit a chondrogenic transcriptomic program(*30*) (high *Col2a1, Acan, Sox9*) with similarities to chondrogenic progenitors in endochondral bone formation and repair. Smad3 has been shown to regulate Sox-factor transcriptional activity in TGFβ-independent manner through physical interactions(*54, 59*). The expansion of the CMC observed also draws an interesting parallel to established findings of accelerated bone and wound healing in *Smad3* knockout mice(*60, 61*). The concomitant increase in *Col2a1* expressing cells in the plaque and increased vascular calcification suggests that this cell type is at least partially responsible for coronary calcifications seen in human coronary artery disease. It remains to be determined whether increased calcification is harmful or protective in terms of plaque rupture risk, since conflicting observational data exists in humans. While increased coronary calcification is correlated with increased risk of myocardial infarction(*62*), local calcification appears to be protective against plaque rupture(*63-65*) and interventions that lower risk of plaque rupture increase calcifications(*66, 67*). Given the multiple populations of SMC-derived transition cells observed in our studies, their relative ratios could possibly determine the quality of calcification as well, which is also considered to confer differential risk of plaque stability.

Recent work by Chen et al.(*46*) has also employed single cell studies to investigate the role of TGFβ signaling in vascular disease, employing a combined Marfan II/ Loeys-Dietz and atherosclerosis mouse model. In keeping with our results reported here, deletion of *Tgfbr2* and *ApoE* genes produced increased plaque burden and consistent with our findings of outward remodeling, led to diffuse aneurysm formation. The mechanism described here by which *Mmp3* expressing cells promote matrix degradation and remodeling were not identified in those studies. While both mouse models showed increased transition of SMC to an endochondral phenotype, there were marked differences related to other SMC transitions. Chen et al. reported an increased inflammatory lesion profile, due in part to SMC transition to the macrophage lineage, but we found no evidence for this type of SMC trans-differentiation. At least one other lab has identified the potential for transition SMC to adopt macrophage marker *Cd68* expression ex vivo(*35*), however an extensive single cell study focused on micro-dissected atherosclerotic lesions failed to find evidence for SMC transition to the macrophage phenotype(*21*). Instead, our studies have validated a specific pro-inflammatory program initiated by SMC, through expression of chemokines that mediate recruitment of this cell type into the plaque. While there were considerable differences in experimental methodology that likely accounts for some of these differences, it also seems likely that modulation of the broad genetic program of TGFβ signaling results in disease features that are distinct from those related to the transcriptional program that is regulated by Smad3, which transduces a portion of the TGFβ signal but in addition receives input from other disease relevant signaling pathways.

Beyond the changes in proportions of the different SMC-derivatives, loss of *Smad3* also resulted in alterations in SMC transition phenotype transcriptomes as a whole. Gene knockout down-regulated several important ECM genes, including *Lox, Mfap5* and *Eln*, whose loss of function mutations have been associated with Mendelian aortopathies. These findings suggest that global transcriptomic changes associated with *Smad3*^*ΔSMC*^ weaken the vascular wall and thus further promote positive or outward vascular remodeling. These finding may also have implications in non-atherosclerotic vasculopathies, such as Marfan and Loeys-Dietz syndromes. Aortopathies such as Marfan’s syndrome have been shown to produce aberrant SMC derived populations that contribute to pathogenesis(*68*). Contrary to our knockout, which decreases TGFβ signaling, collagen vascular dilated aortopathies are thought to be associated with an increase in TGFβ signaling(*69*). Specific pathogenic mutations in the Smad3 MH2 domain have been linked to Loeys-Dietz syndrome 3 in humans with evidence pointing to both TGFβ gain of function(*70*) and loss of function(*71, 72*) mutations. These contradictory findings suggest a potential decoupling of Smad3 function in vascular SMC from its role as a TGFβ signaling secondary messenger. Given the role of the MH2 domain in protein-protein interaction and our finding of the importance of the transcriptional network regulated by Smad3-dependent activation of Sox and Hox -regulated transcriptional programs, our work could offer a unifying hypothesis whereby the aortic dilatation phenotype is driven by loss of downstream interactions of Smad3. Augmentation of the Sox and Hox transcriptional program to boost expression of these critical ECM genes and possibly change SMC-derived populations in the vascular wall may therefore offer therapeutic implications for some aortopathies and warrants further investigation.

The limitation of our lineage tracing methodology does not allow us to definitively determine the plasticity of SMC transition phenotypes and their ability to shuttle between the different populations. It is unclear if CMCs or the new *Mmp3*-expressing remodeling R-SMC can be induced to adopt the phenotype of each other or return to the FMC phenotype. Nor can we be certain that a specific cell can differentiate only into CMC or R-SMC, or whether they are distinct populations at the time of their activation from mature SMC. Though the lineage inference using pseudo-time techniques presented here offers evidence that their common progenitor lies in the FMC stage, due to the limitations of these techniques, additional lineage labeling experiment with cre-recombinases specific for each of these populations would be required to directly address these possibilities.

Overall, this study extends existing literature identifying smooth muscle transition cell phenotype as a primary activator of lipid-independent genetic risk for coronary artery disease and affirms the ability of GWAS to identify critical cellular and molecular pathways involved in the pathology of atherosclerosis.

## Methods

### Mouse strains

SMC-specific lineage tracing and *Smad3* knockout was generated by a well-characterized BAC transgene that expresses a tamoxifen-inducible Cre recombinase driven by the SMC-specific *Myh11* promoter (*Tg*^Myh11-CreERT2^; 019079; JAX). These mice were bred with a floxed-stop-flox *tdTomato* fluorescent reporter line (B6.Cg-*Gt(ROSA)26Sor*^*tm14(CAGtdTomato)Hze*^/J; 007914; JAX) to allow SMC-specific lineage tracing. *Smad3* conditional knockout were obtained from Matzuk lab from UTSW (*41, 42*)with LoxP sites flanking exons 2 and 3 which contains Smad3 DNA binding domain and creates a non-functioning frame-shift mutation after deletion (Zhu et al, cell 1998). All mice were back-crossed onto the C56BL/6 *ApoE*^*−/−*^ background. As the Cre-expressing BAC was integrated into the Y chromosome, all lineage-tracing mice in the study were male. The animal study protocol was approved by the Administrative Panel on Laboratory Animal Care at Stanford University.

### Induction of lineage marker and *Smad3* knockout by Cre recombinase

For all experiments, tamoxifen gavage schedule was as follows: two doses of tamoxifen, at 0.2□mg□g^−1^ bodyweight, were administered by oral gavage at 7-8□weeks of age, with each dose separated by 72-96□hrs. HFD was started (101511; Dyets; 21% anhydrous milk fat, 19% casein and 0.15% cholesterol) after the second gavage.

### Mouse aortic root/ascending aorta cell dissociation

Immediately after sacrifice, mice were perfused with phosphate buffered saline (PBS). The aortic root and ascending aorta were excised, up to the level of the brachiocephalic artery. Tissue was washed three times in PBS, placed into an enzymatic dissociation cocktail (2□U□ml^−1^ Liberase (5401127001; Sigma–Aldrich) and 2□U□ml^−1^ elastase (LS002279; Worthington) in Hank’s Balanced Salt Solution (HBSS)) and minced. After incubation at 37□°C for 1□h, the cell suspension was strained and then pelleted by centrifugation at 500*g* for 5□min. The enzyme solution was then discarded, and cells were resuspended in fresh HBSS. To increase biological replication, multiple mice were used to obtain single-cell suspensions at each time point. For each scRNA capture, 2 mice were used. 4 separate pairs of isolation were performed for control and *Smad3*^*ΔSMC*^, but one control 10X capture unexpectedly failed resulting in a final of 3 captures of control and 4 captures from conditional KO that was included in the analysis. Cells were sorted FACS sorted based off tdTomato expression. *tdT*^+^ cells (considered to be of SMC lineage) and *tdT*^−^ cells were then captured on separate but parallel runs of the same scRNA-Seq workflow(gating strategy and threshold identical to those published in previous work by Wirka et al^24^), and datasets were later combined for all subsequent analyses.

### Single-cell capture and library preparation and sequencing

All single-cell capture and library preparation was performed at the Stanford Functional Genomics Facility and Stanford Genomic Sequencing Service Center. Cells were loaded into a 10x Genomics microfluidics chip and encapsulated with barcoded oligo-dT-containing gel beads using the 10x Genomics Chromium controller according to the manufacturer’s instructions. Single-cell libraries were then constructed according to the manufacturer’s instructions. Libraries from individual samples were multiplexed into one lane before sequencing on an Illumina platforms with targeted depth of 50,000 reads per cell. At this depth, sequencing saturation was ∼25-35%.

### Human coronary artery cell section

Human coronary arteries used in this study were dissected from explanted hearts of transplant recipients, and were obtained from the Human Biorepository Tissue Research Bank under the Department of Cardiothoracic Surgery from consenting patients with approval from the Stanford University Institutional Review Board as previously described.

### Preparation of mouse aortic root sections

Immediately after sacrifice, mice were perfused with 0.4% paraformaldehyde (PFA). The mouse aortic root and proximal ascending aorta, along with the base of the heart, was excised and immersed in 4% PFA at 4□°C for 24□hrs. After passing through a sucrose gradient, tissue was frozen in optimal cutting temperature compound (OCT) to make blocks. Blocks were cut into 7-µm-thick sections for further analysis.

### Immunohistochemistry

Slides were air dried and OCT was removed with two washes in deionized water. Slides were immersed in 4% PFA for 2□min, followed by four washes in deionized water. Slides were dried and sections were encircled with a liquid-blocking pen, followed by Peroxidazed (PX968; Biocare Medical) treatment for 5□min. Sections were washed three times with deionized water, and then incubated with PBS with 5% BSA and 5% FBS or Rodent Block M reagent (RBM961; Biocare Medical) for 60□min.

Sections were washed twice in Tris-buffered saline (TBS), then incubated overnight at 4□°C with an anti-SM22alpha rabbit polyclonal primary antibody (1:300 dilution; ab14106; Abcam), a Mmp3 Rabbit monoclonal antibody (1:200 dilution; Abcam 52915) or a CD68 rabbit polyclonal antibody (1:400 dilution; ab125212; Abcam). Sections were washed for 5□min twice with TBS and then incubated with the Rabbit-on-Rodent HRP Polymer (RMR622; Biocare Medical) for 30□min at room temperature.

Sections were washed twice with TBS and then incubated with the Betazoid DAB chromogen reagents (BDB2004; Biocare Medical) for 4□min at room temperature followed by mounting with EcoMount medium (EM897L; Biocare Medical). The processed sections were visualized using a Leica DM5500 microscope objective magnifications, and images were obtained using Leica Application Suite X software. Sections obtained at equal distance measured from the superior margin of the aortic sinus were used for comparison. Areas of interest were quantified using ImageJ (National Institutes of Health) software, and compared using a two-sided *t*-test. Lesion size were defined by the area encompassing the intimal edge of the lesion to the border of tagln positive intima-media junction. Area encompassed by the vessel media was defined by area encircled by the outer edge of Tagln staining of vessel media. All area quantification was performed in a genotype blinded fashion with image J using length information embedded in exported files. Von Kossa stain was performed using Abcam 150687 kit with manufacturer’s recommended protocol with 90 minute development time. All biological replicates for each staining were performed simultaneously on position-matched aortic root sections to limit intra-experimental variance. Folded sections that were uninterpretable after processing were removed.

### RNAscope assay

Slides were processed according to the manufacturer’s instructions, and all reagents were obtained from ACD Bio. In short, slides were washed once in PBS, then immersed in 1× Target Retrieval reagent at 100□°C for 5□min. Slides were washed twice in deionized water, immersed in 100% ethanol and air dried, and sections were encircled with a liquid-blocking pen. Sections were incubated with Protease III reagent for 30□min at 40□°C, then washed twice with deionized water. Sections were incubated with commercially available probes against mouse Mmp3, Col2a1, and human *MMP3* or a negative control probe for 2□hrs at 40□°C. Colorimetric assays were performed per the manufacturer’s instructions.

### Analysis of scRNA-Seq data

Fastq files from each experimental time point and mouse genotype were aligned to the reference genome (mm10) individually using CellRanger Software (10x Genomics). Individual datasets were aggregated using the CellRanger aggr command without subsampling normalization. The aggregated dataset was then analyzed using the R package Seurat(*73*). The dataset was trimmed of cells expressing fewer than 750 genes, and genes expressed in fewer than 50 cells. The number of genes, number of unique molecular identifiers and percentage of mitochondrial genes were examined to identify outliers. As an unusually high number of genes can result from a ‘doublet’ event, in which two different cell types are captured together with the same barcoded bead, cells with >6000 genes were discarded. Cells containing >7.5% mitochondrial genes were presumed to be of poor quality and were also discarded. The gene expression values then underwent library-size normalization and normalized using established Single-Cell-Transform function built into Seurat. Principal component analysis was used for dimensionality reduction, followed by clustering in principal component analysis space using a graph-based clustering approach via Louvain algorithm UMAP was then used for two-dimensional visualization of the resulting clusters. Lineage inference was performed using Slingshot with available Slingshot software in R using converted Seurat object into singlecellexperiment objects. Analysis, visualization and quantification of gene expression and generation of gene module scores were performed using Seurat’s built in function such as “FeaturePlot”, “VlnPlot”, “AddModuleScore”, and “FindMarker.” List of genes associated with each GO category was pulled from Geneontology.org. Panther / DAVID / GO / GREAT analysis was performed using web-based platform at Geneontology.org, Great.Stanford.Edu, and David.ncifcrf.gov. Top 1000 genes expressed in modulated SMC was defined by the highest expressing 1000 transcripts (based off average expression) from scRNA data in all de-differentiated lineage traced cells. Promoter/5’-Regulatory region of genes were extracted utilizing UCSC table browser based off 1kb upstream of TSS of transcripts. Motif analysis was performed using freely available HOMER software(*74*) with findMotifGenome function. The regulatory region of the top 1000 gene was used as the background as bases for motif enrichment.

### HCASMC culture/experiments

Cells were cultured in smooth muscle growth medium (Lonza; catalog number: CC-3182) supplemented with human epidermal growth factor, insulin, human basic fibroblast growth factor and 5% FBS, according to the manufacturer’s instructions. All HCASMC lines were used at passages 4–8. siRNA knockdown were performed using Lipofectamine RNAiMax (Life Technologies) using manufacturer’s recommended protocol at 50pg siRNA / 100,000 cells. Cells were allowed to recover in SMC growth medium (with or without additional growth factor) for 36 hours prior to RNA harvest. Recombinant TGFb (PeproTech 100-21) concentration used in stimulation was 10ng/ml.

### Co-IP Experiment

Myc-Flag-tagged SOX9 (PS100016 Origene) and 6x-HIS-tagged-HOXB2 (Addgene 8522) were obtained from commercial vendors and cloned into pCMV6 vector and transfected into HEK cells. The cells were allowed to recover for 36 hours after media change and Nuclear-Complex Co-IP was performed using commercially available Nuclear-Complex Co-IP kit from ActiveMotif(54001) with manufacturer’s recommended protocol using mouse Anti-Flag (Sigma F3165) or mouse Anti-His (Abcam 18184) antibody for immunoprecipitation, followed by blotting using Rabbit anti-Smad3 antibody (Cell Signaling 9523S), followed by Anti-Rabbit HRP (Cell Signaling 7074S) and detected via Luminata Forte Western HRP substrate (Millipore).

### Luciferase experiments

For Luciferase experiments, evolutionary conserved region of human MFAP5 regulatory region (chr12:8815212-8815569) was cloned from human genomic DNA and placed into pLuc-MCS vector, whereas inert/scramble similar length spacer was cloned into baseline pLuc-MCS as control. pCMV6-empty, and cloned pCMV6-Flag-SOX9 or pCMV6-his-HOXB2 were transfected into cells via lipofectamine 2000 along with respective luciferase and Renilla vector. Media was changed after 6□h, and dual-luciferase activity (Promega) was recorded after 24□h using a SpectraMax L luminometer (Molecular Devices). Relative luciferase activity (firefly/*Renilla* luciferase ratio) is expressed as the fold change over control conditions.

### TUNEL Assay

TUNEL assay was performed on cryopreserved aortic root sections using commercially available chromogenic TUNEL assay kits. Quantification was performed on 40X magnification at one random point on each cusp and TUNEL+ nuclei was counted manually in a blinded manner.

### Statistical methods

Differentially expressed genes in the scRNA-Seq data were identified using a Wilcoxon rank-sum test. Distribution of cells within defined-populations was tested via X-square test.

Significance determination of histological measurement, luciferase studies, qPCR results, and composite gene-score were done via two-tailed T-test. Multiple comparisons were corrected via Bonferroni correction when necessary.

## Acknowledgements

Special thanks to the Krista Hennig, Yana Ryan, Peter Mcguire and Hassan Chiab at the Stanford Genomic Sequencing and Service Center (GSSC) for performing 10x capture, library construction, and sequencing. Stanford shared FACS facility for required FACS analysis and experiments. Thanks to the Matzuk lab for providing us with conditional Smad3 knockout mice. Illustrations were made with BioRender software.

## Supplemental Figure Legends

**Sup Fig 1:**
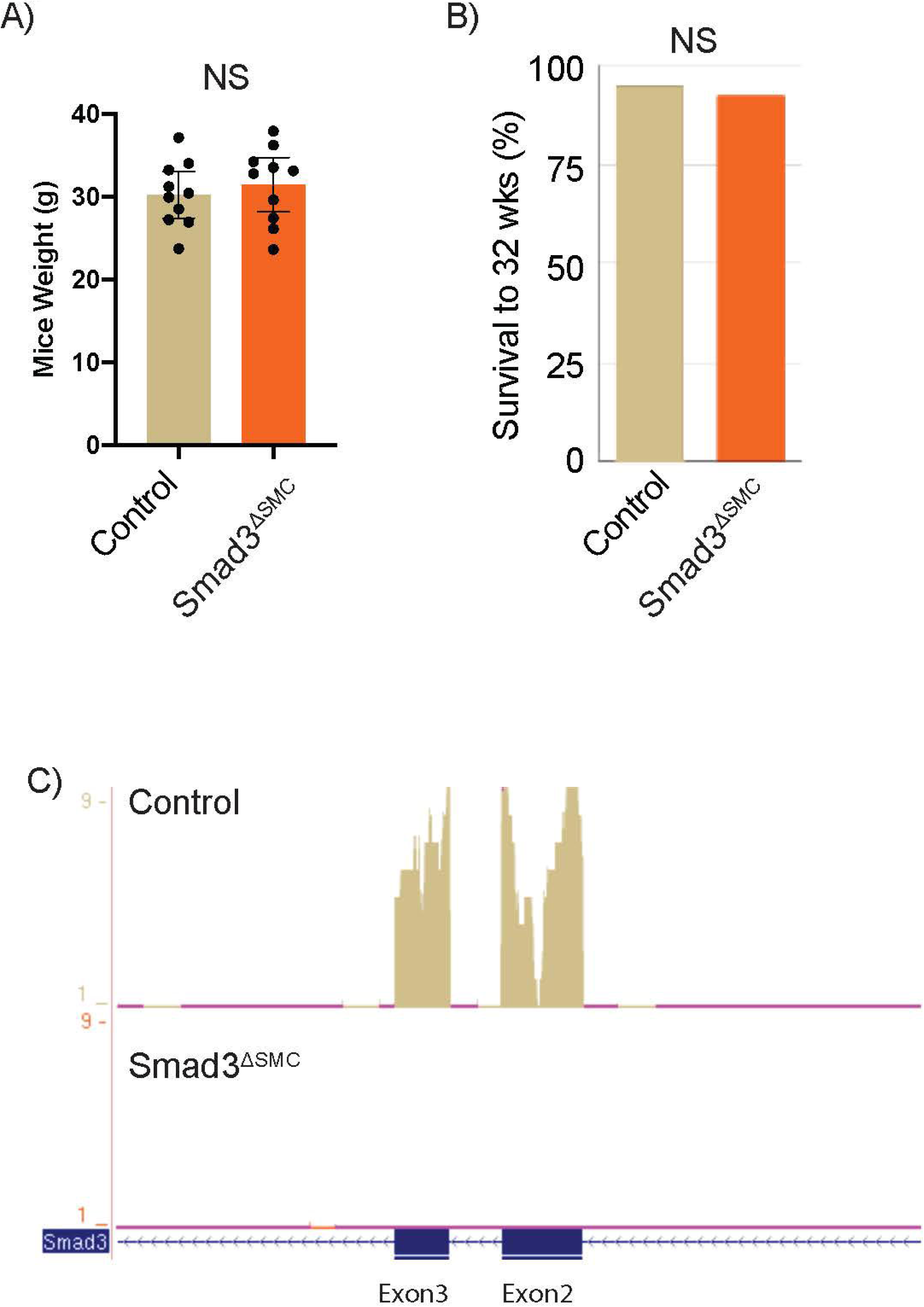
(A) Measured weight of experimental mice used in sections at time of sacrifice. (B) Percent of total experimental mice cohort that survived to final time point at 32 weeks. (C) Aligned captured mRNA sequence of tdTomato positive cells in control (top) and *Smad3*^*ΔSMC*^ tdTomato cells, showing absence of reads mapping to exon 2-3 which are flanked by LoxP sites.

**Sup Fig 2:**
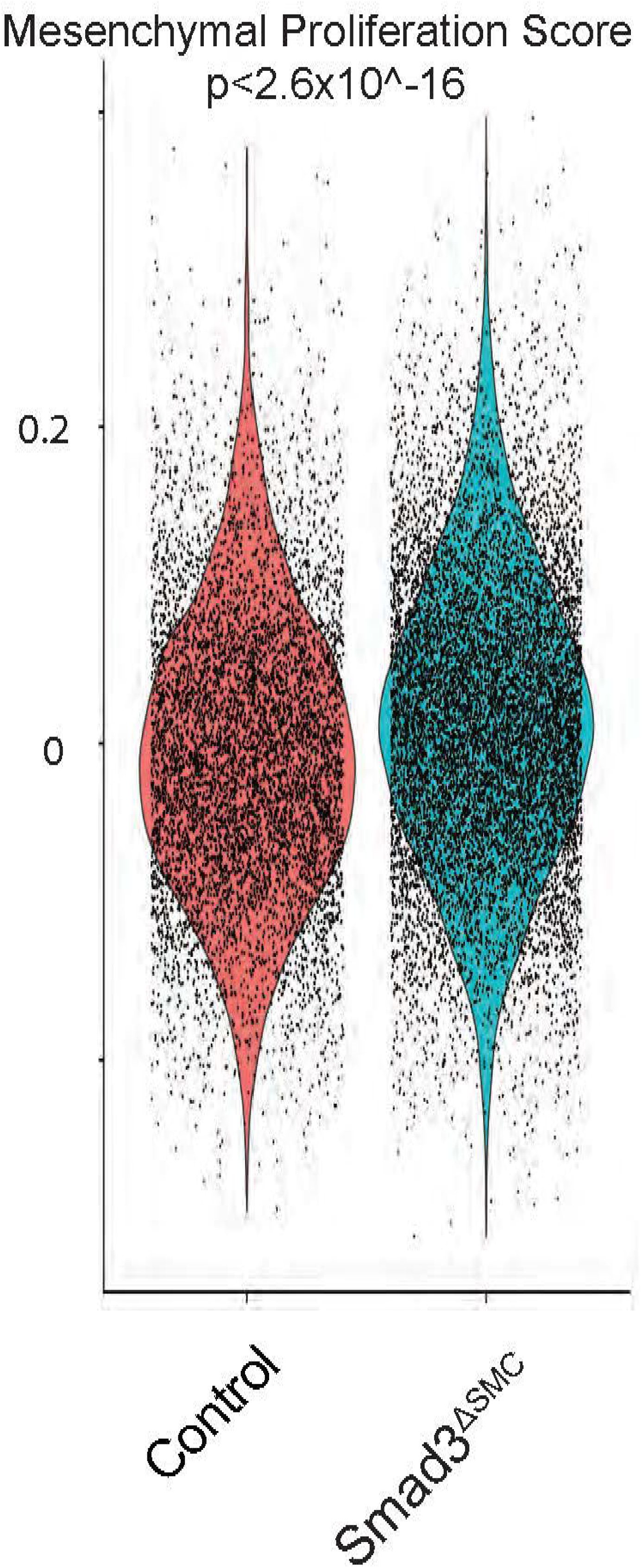
Mesenchymal proliferation score of de-differentiated SMC in control and SMC specific *Smad3*^*ΔSMC*^ mice.

**Sup Fig 3:**
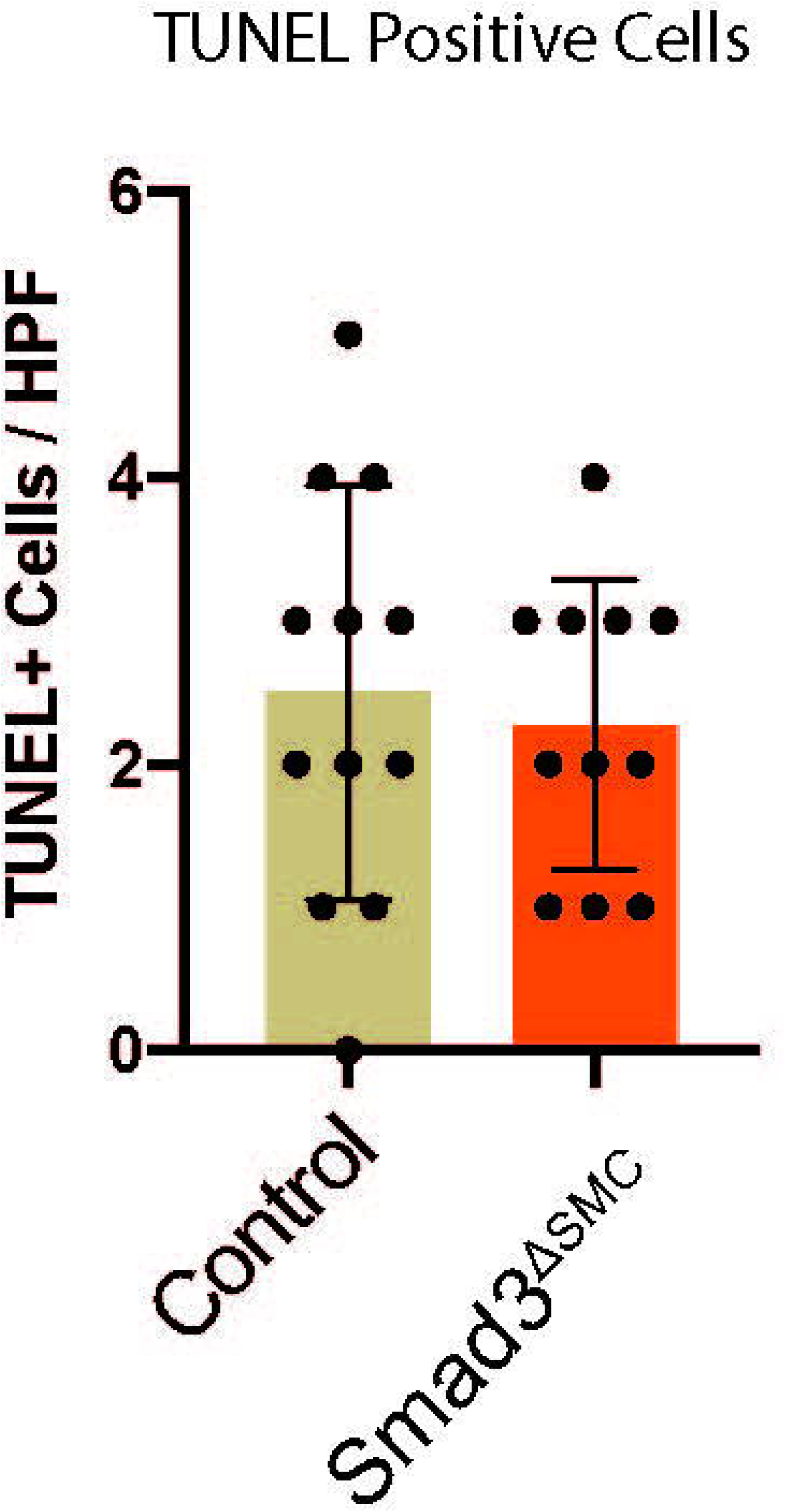
Number of TUNEL labeled apoptotic cells per high power fiedl (HPF) in control and *Smad3*^*ΔSMC*^ mice.

**Sup Fig 4:**
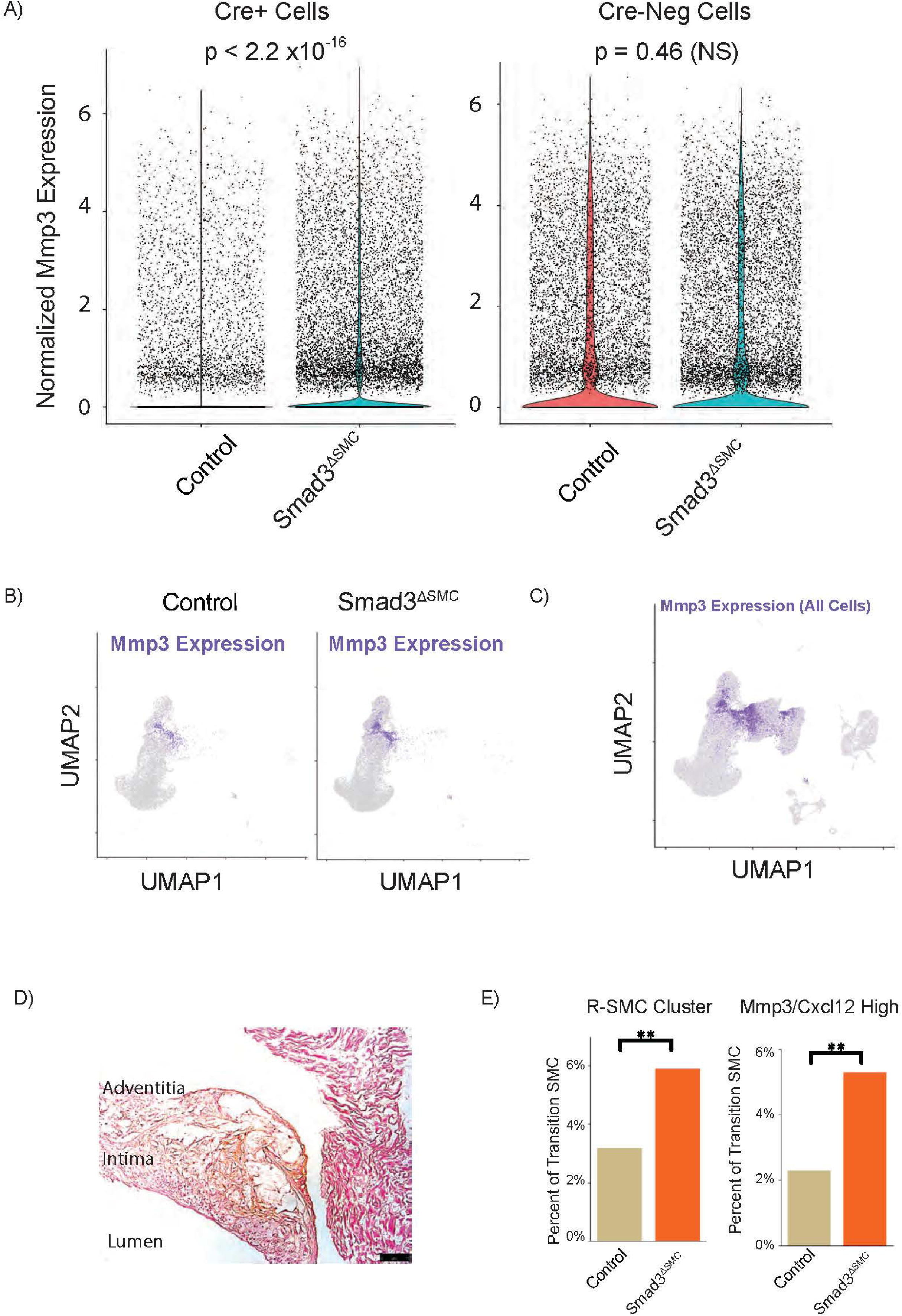
(A) Expression of *Mmp3* in lineage labeled (Cre positive) and non-lineage labelled (Cre negative) cells based on scRNAseq data. (B) Featureplot of lineage-traced cells expressing *Mmp3* in control and *Smad3*^*ΔSMC*^ aortic root. C) Featureplot of *Mmp3* expression in all cells. D) High power view of section of atherosclerotic plaque with region of broken elastic lamina stained for Mmp3 expression (orange-brown color). E) Fraction of transition SMC in control and *Smad3*^*ΔSMC*^ lesions with R-SMC fate as defined by unbiased clustering (left) and by concurrent high Mmp3/CxCl12 expression (right).

**Sup Fig 5:**
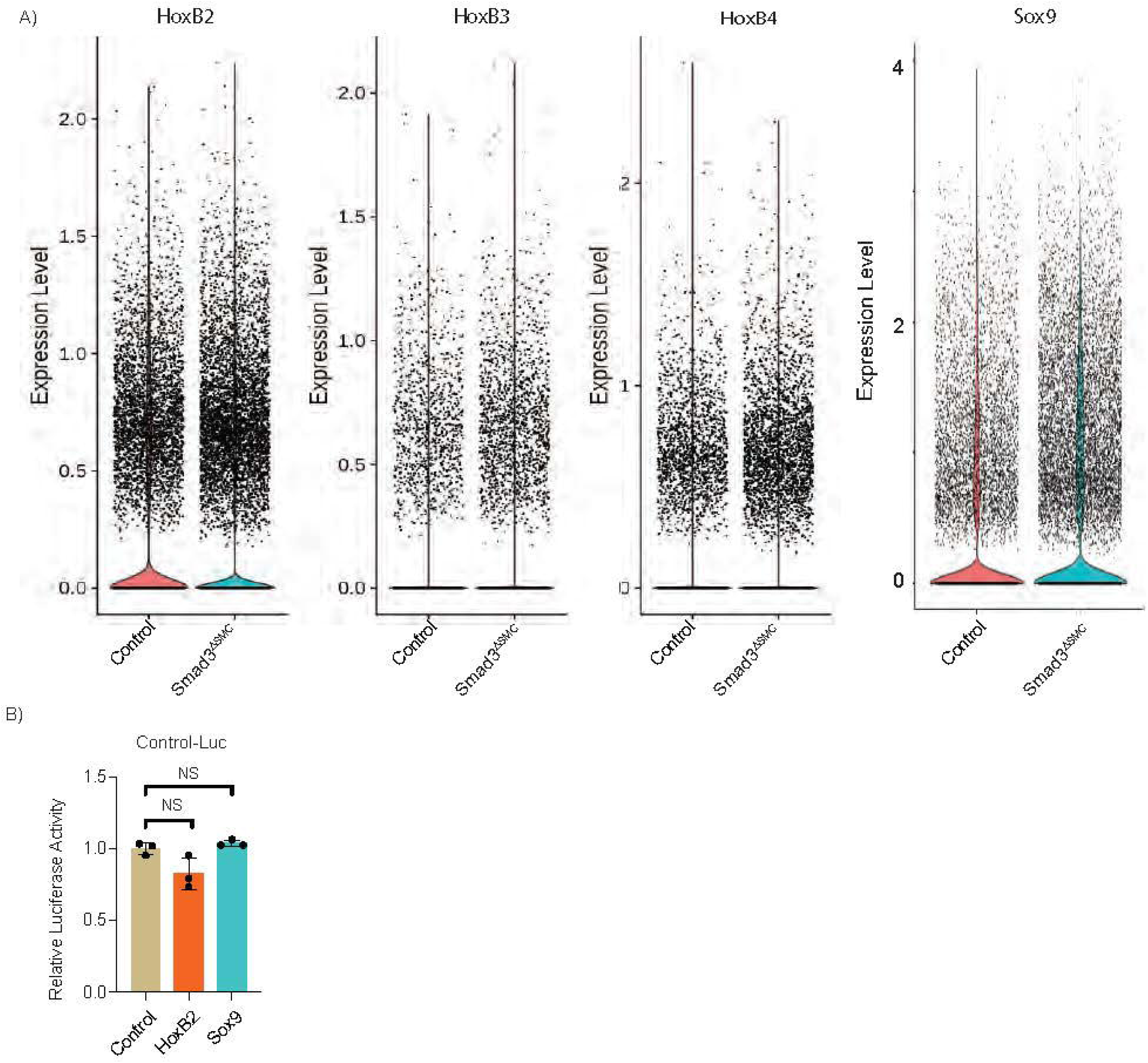
Individual cell expression of *HoxB2, HoxB3, HoxB4*, and *Sox9* in lineage labeled cells in control and *Smad3*^*ΔSMC*^ aortic root.

**Sup Fig 6:**
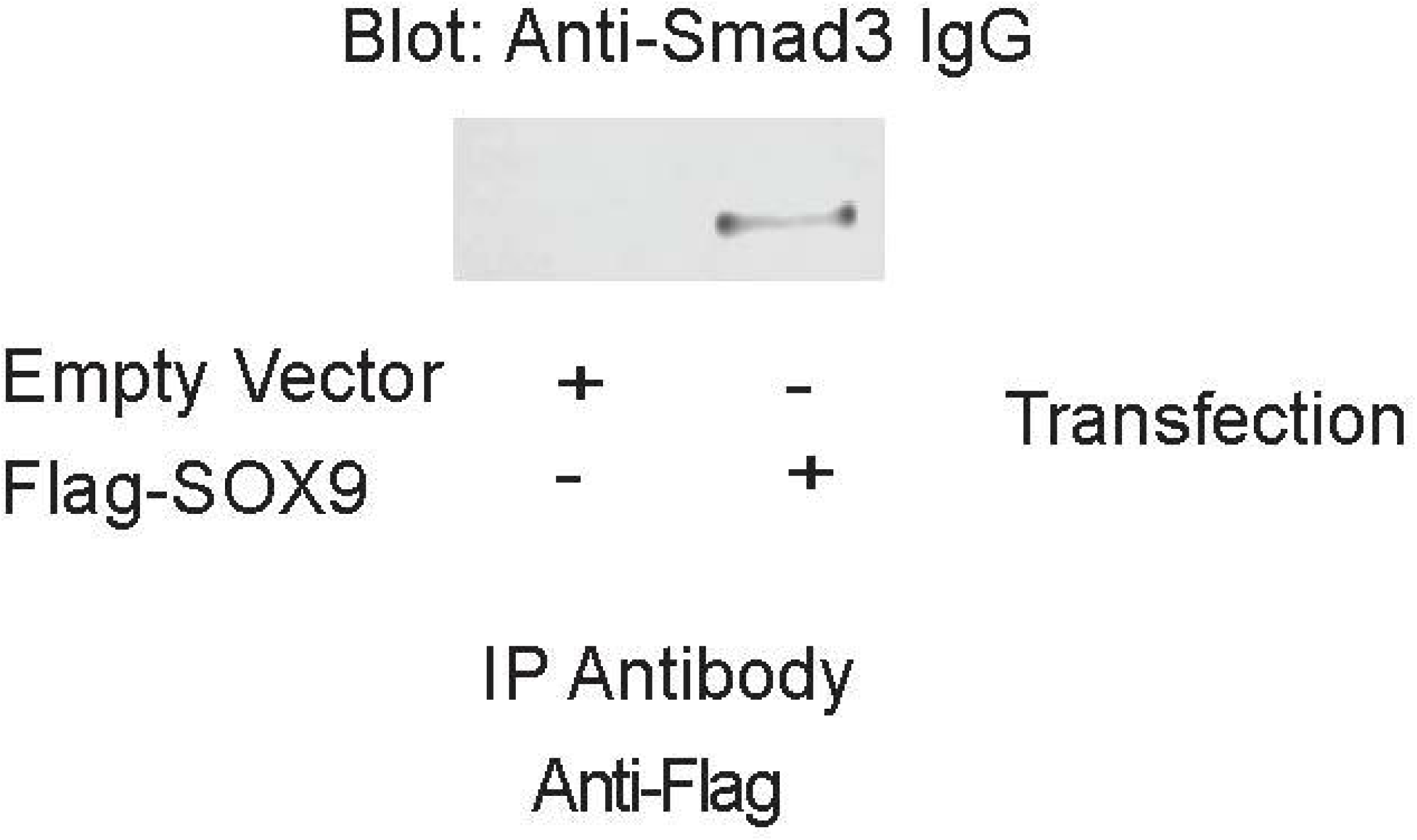
Additional replicates of Flag-SOX9 with endogenous SMAD3 IP with additional controls in anti-flag antibody.

## Supplemental Files

Supplementary Table 1: Differentially expressed gene in R-SMC vs other transition SMC.

Supplementary Table 2: Differentially expressed gene in transition SMC in *Smad3*^*ΔSMC*^ vs control.

## Data Availability Statement

The data generated during this study will be available from the corresponding author upon reasonable request during review, and will be uploaded to GEO database after acceptance of the manuscript.

## References

1. CDC. (2020).

2. K. Reynolds et al., Trends in Incidence of Hospitalized Acute Myocardial Infarction in the Cardiovascular Research Network (CVRN). Am J Med 130, 317–327 (2017).

3. S. Sidney et al., Recent Trends in Cardiovascular Mortality in the United States and Public Health Goals. JAMA Cardiol 1, 594–599 (2016).

4. M. S. Sabatine, S. M. Wasserman, E. A. Stein, PCSK9 Inhibitors and Cardiovascular Events. N Engl J Med 373, 774–775 (2015).

5. P. M. Ridker et al., Cardiovascular Efficacy and Safety of Bococizumab in High-Risk Patients. N Engl J Med 376, 1527–1539 (2017).

6. P. M. Ridker et al., Antiinflammatory Therapy with Canakinumab for Atherosclerotic Disease. N Engl J Med 377, 1119–1131 (2017).

7. P. M. Ridker et al., Low-Dose Methotrexate for the Prevention of Atherosclerotic Events. N Engl J Med 380, 752–762 (2019).

8. R. A. Harrington, Targeting Inflammation in Coronary Artery Disease. N Engl J Med 377, 1197–1198 (2017).

9. M. Nikpay et al., A comprehensive 1,000 Genomes-based genome-wide association meta-analysis of coronary artery disease. Nat Genet 47, 1121–1130 (2015).

10. P. van der Harst, N. Verweij, Identification of 64 Novel Genetic Loci Provides an Expanded View on the Genetic Architecture of Coronary Artery Disease. Circ Res 122, 433–443 (2018).

11. I. Braenne et al., Prediction of Causal Candidate Genes in Coronary Artery Disease Loci. Arterioscler Thromb Vasc Biol 35, 2207–2217 (2015).

12. C. L. Miller, M. Pjanic, T. Quertermous, From Locus Association to Mechanism of Gene Causality: The Devil Is in the Details. Arterioscler Thromb Vasc Biol 35, 2079–2080 (2015).

13. B. Liu et al., Genetic Regulatory Mechanisms of Smooth Muscle Cells Map to Coronary Artery Disease Risk Loci. Am J Hum Genet 103, 377–388 (2018).

14. K. Sakakura et al., Pathophysiology of atherosclerosis plaque progression. Heart Lung Circ 22, 399–411 (2013).

15. P. K. Shah, Mechanisms of plaque vulnerability and rupture. J Am Coll Cardiol 41, 15S–22S (2003).

16. E. Falk, M. Nakano, J. F. Bentzon, A. V. Finn, R. Virmani, Update on acute coronary syndromes: the pathologists’ view. Eur Heart J 34, 719–728 (2013).

17. M. Ferencik et al., Use of High-Risk Coronary Atherosclerotic Plaque Detection for Risk Stratification of Patients With Stable Chest Pain: A Secondary Analysis of the PROMISE Randomized Clinical Trial. JAMA Cardiol 3, 144–152 (2018).

18. S. B. Puchner et al., High-risk plaque detected on coronary CT angiography predicts acute coronary syndromes independent of significant stenosis in acute chest pain: results from the ROMICAT-II trial. J Am Coll Cardiol 64, 684–692 (2014).

19. M. R. Ward, G. Pasterkamp, A. C. Yeung, C. Borst, Arterial remodeling. Mechanisms and clinical implications. Circulation 102, 1186–1191 (2000).

20. M. C. Williams et al., Coronary Artery Plaque Characteristics Associated With Adverse Outcomes in the SCOT-HEART Study. J Am Coll Cardiol 73, 291–301 (2019).

21. G. F. Alencar et al., The Stem Cell Pluripotency Genes Klf4 and Oct4 Regulate Complex SMC Phenotypic Changes Critical in Late-Stage Atherosclerotic Lesion Pathogenesis. Circulation, (2020).

22. C. L. Miller et al., Integrative functional genomics identifies regulatory mechanisms at coronary artery disease loci. Nat Commun 7, 12092 (2016).

23. S. T. Nurnberg et al., Coronary Artery Disease Associated Transcription Factor TCF21 Regulates Smooth Muscle Precursor Cells that Contribute to the Fibrous Cap. Genom Data 5, 36–37 (2015).

24. L. S. Shankman et al., KLF4-dependent phenotypic modulation of smooth muscle cells has a key role in atherosclerotic plaque pathogenesis. Nat Med 21, 628–637 (2015).

25. R. C. Wirka et al., Atheroprotective roles of smooth muscle cell phenotypic modulation and the TCF21 disease gene as revealed by single-cell analysis. Nat Med 25, 1280–1289 (2019).

26. J. Chappell et al., Extensive Proliferation of a Subset of Differentiated, yet Plastic, Medial Vascular Smooth Muscle Cells Contributes to Neointimal Formation in Mouse Injury and Atherosclerosis Models. Circ Res 119, 1313–1323 (2016).

27. K. Jacobsen et al., Diverse cellular architecture of atherosclerotic plaque derives from clonal expansion of a few medial SMCs. JCI Insight 2, (2017).

28. A. Misra et al., Integrin beta3 regulates clonality and fate of smooth muscle-derived atherosclerotic plaque cells. Nat Commun 9, 2073 (2018).

29. C. E. Murry, C. T. Gipaya, T. Bartosek, E. P. Benditt, S. M. Schwartz, Monoclonality of smooth muscle cells in human atherosclerosis. Am J Pathol 151, 697–705 (1997).

30. J. B. Kim et al., Environment-Sensing Aryl Hydrocarbon Receptor Inhibits the Chondrogenic Fate of Modulated Smooth Muscle Cells in Atherosclerotic Lesions. Circulation 142, 575–590 (2020).

31. D. Iyer et al., Coronary artery disease genes SMAD3 and TCF21 promote opposing interactive genetic programs that regulate smooth muscle cell differentiation and disease risk. PLoS Genet 14, e1007681 (2018).

32. J. B. Kim et al., TCF21 and the environmental sensor aryl-hydrocarbon receptor cooperate to activate a pro-inflammatory gene expression program in coronary artery smooth muscle cells. PLoS Genet 13, e1006750 (2017).

33. C. L. Miller et al., Disease-related growth factor and embryonic signaling pathways modulate an enhancer of TCF21 expression at the 6q23.2 coronary heart disease locus. PLoS Genet 9, e1003652 (2013).

34. C. L. Miller et al., Coronary heart disease-associated variation in TCF21 disrupts a miR-224 binding site and miRNA-mediated regulation. PLoS Genet 10, e1004263 (2014).

35. H. Pan et al., Single-Cell Genomics Reveals a Novel Cell State During Smooth Muscle Cell Phenotypic Switching and Potential Therapeutic Targets for Atherosclerosis in Mouse and Human. Circulation, (2020).

36. D. J. Grainger, Transforming growth factor beta and atherosclerosis: so far, so good for the protective cytokine hypothesis. Arterioscler Thromb Vasc Biol 24, 399–404 (2004).

37. I. Toma, T. A. McCaffrey, Transforming growth factor-beta and atherosclerosis: interwoven atherogenic and atheroprotective aspects. Cell Tissue Res 347, 155–175 (2012).

38. Y. Shi et al., Crystal structure of a Smad MH1 domain bound to DNA: insights on DNA binding in TGF-beta signaling. Cell 94, 585–594 (1998).

39. J. Massague, S. W. Blain, R. S. Lo, TGFbeta signaling in growth control, cancer, and heritable disorders. Cell 103, 295–309 (2000).

40. M. Morikawa, R. Derynck, K. Miyazono, TGF-beta and the TGF-beta Family: Context-Dependent Roles in Cell and Tissue Physiology. Cold Spring Harb Perspect Biol 8, (2016).

41. M. Kriseman et al., Uterine double-conditional inactivation of Smad2 and Smad3 in mice causes endometrial dysregulation, infertility, and uterine cancer. Proc Natl Acad Sci U S A 116, 3873–3882 (2019).

42. Q. Li et al., Redundant roles of SMAD2 and SMAD3 in ovarian granulosa cells in vivo. Mol Cell Biol 28, 7001–7011 (2008).

43. Y. Zhu, J. A. Richardson, L. F. Parada, J. M. Graff, Smad3 mutant mice develop metastatic colorectal cancer. Cell 94, 703–714 (1998).

44. Y. K. Hong et al., Prox1 is a master control gene in the program specifying lymphatic endothelial cell fate. Dev Dyn 225, 351–357 (2002).

45. J. Wilting et al., The transcription factor Prox1 is a marker for lymphatic endothelial cells in normal and diseased human tissues. FASEB J 16, 1271–1273 (2002).

46. P. Y. Chen et al., Smooth Muscle Cell Reprogramming in Aortic Aneurysms. Cell Stem Cell 26, 542–557 e511 (2020).

47. I. Tirosh et al., Dissecting the multicellular ecosystem of metastatic melanoma by single-cell RNA-seq. Science 352, 189–196 (2016).

48. K. Street et al., Slingshot: cell lineage and pseudotime inference for single-cell transcriptomics. BMC Genomics 19, 477 (2018).

49. M. R. Alexander et al., Genetic inactivation of IL-1 signaling enhances atherosclerotic plaque instability and reduces outward vessel remodeling in advanced atherosclerosis in mice. J Clin Invest 122, 70–79 (2012).

50. C. Y. McLean et al., GREAT improves functional interpretation of cis-regulatory regions. Nat Biotechnol 28, 495–501 (2010).

51. M. Barbier et al., MFAP5 loss-of-function mutations underscore the involvement of matrix alteration in the pathogenesis of familial thoracic aortic aneurysms and dissections. Am J Hum Genet 95, 736–743 (2014).

52. D. C. Guo et al., LOX Mutations Predispose to Thoracic Aortic Aneurysms and Dissections. Circ Res 118, 928–934 (2016).

53. A. Pinard, G. T. Jones, D. M. Milewicz, Genetics of Thoracic and Abdominal Aortic Diseases. Circ Res 124, 588–606 (2019).

54. Furumatsu, M. Tsuda, N. Taniguchi, Y. Tajima, H. Asahara, Smad3 induces chondrogenesis through the activation of SOX9 via CREB-binding protein/p300 recruitment. J Biol Chem 280, 8343–8350 (2005).

55. M. W. Majesky et al., Differentiated Smooth Muscle Cells Generate a Subpopulation of Resident Vascular Progenitor Cells in the Adventitia Regulated by Klf4. Circ Res 120, 296–311 (2017).

56. S. Beyzade et al., Influences of matrix metalloproteinase-3 gene variation on extent of coronary atherosclerosis and risk of myocardial infarction. J Am Coll Cardiol 41, 2130–2137 (2003).

57. G. B. Bulut et al., KLF4 (Kruppel-Like Factor 4)-Dependent Perivascular Plasticity Contributes to Adipose Tissue inflammation. Arterioscler Thromb Vasc Biol, ATVBAHA120314703 (2020).

58. R. A. Nemenoff et al., SDF-1alpha induction in mature smooth muscle cells by inactivation of PTEN is a critical mediator of exacerbated injury-induced neointima formation. Arterioscler Thromb Vasc Biol 31, 1300–1308 (2011).

59. S. J. Vervoort et al., SOX4 can redirect TGF-beta-mediated SMAD3-transcriptional output in a context-dependent manner to promote tumorigenesis. Nucleic Acids Res 46, 9578–9590 (2018).

60. G. S. Ashcroft et al., Role of Smad3 in the hormonal modulation of in vivo wound healing responses. Wound Repair Regen 11, 468–473 (2003).

61. G. S. Ashcroft et al., Mice lacking Smad3 show accelerated wound healing and an impaired local inflammatory response. Nat Cell Biol 1, 260–266 (1999).

62. R. L. McClelland et al., 10-Year Coronary Heart Disease Risk Prediction Using Coronary Artery Calcium and Traditional Risk Factors: Derivation in the MESA (Multi-Ethnic 2. Study of Atherosclerosis) With Validation in the HNR (Heinz Nixdorf Recall) Study and the DHS (Dallas Heart Study). J Am Coll Cardiol 66, 1643–1653 (2015).

63. M. H. Criqui et al., Calcium density of coronary artery plaque and risk of incident cardiovascular events. JAMA 311, 271–278 (2014).

64. S. Motoyama et al., Multislice computed tomographic characteristics of coronary lesions in acute coronary syndromes. J Am Coll Cardiol 50, 319–326 (2007).

65. S. J. Nicholls et al., Coronary artery calcification and changes in atheroma burden in response to established medical therapies. J Am Coll Cardiol 49, 263–270 (2007).

66. V. L. Aengevaeren et al., Relationship Between Lifelong Exercise Volume and Coronary Atherosclerosis in Athletes. Circulation 136, 138–148 (2017).

67. R. Puri et al., Impact of statins on serial coronary calcification during atheroma progression and regression. J Am Coll Cardiol 65, 1273–1282 (2015).

68. A. J. Pedroza et al., Single-Cell Transcriptomic Profiling of Vascular Smooth Muscle Cell Phenotype Modulation in Marfan Syndrome Aortic Aneurysm. Arterioscler Thromb Vasc Biol 40, 2195–2211 (2020).

69. P. Matt et al., Circulating transforming growth factor-beta in Marfan syndrome. Circulation 120, 526–532 (2009).

70. I. M. van de Laar et al., Mutations in SMAD3 cause a syndromic form of aortic aneurysms and dissections with early-onset osteoarthritis. Nat Genet 43, 121–126 (2011).

71. E. S. Regalado et al., Exome sequencing identifies SMAD3 mutations as a cause of familial thoracic aortic aneurysm and dissection with intracranial and other arterial aneurysms. Circ Res 109, 680–686 (2011).

72. I. M. van de Laar et al., Phenotypic spectrum of the SMAD3-related aneurysms-osteoarthritis syndrome. J Med Genet 49, 47–57 (2012).

73. Stuart et al., Comprehensive Integration of Single-Cell Data. Cell 177, 1888–1902 e1821 (2019).

74. S. Heinz et al., Simple combinations of lineage-determining transcription factors prime cis-regulatory elements required for macrophage and B cell identities. Mol Cell 38, 576–589 (2010).

